# Reverse-complement parameter sharing improves deep learning models for genomics

**DOI:** 10.1101/103663

**Authors:** Avanti Shrikumar, Peyton Greenside, Anshul Kundaje

## Abstract

Deep learning approaches that have produced breakthrough predictive models in computer vision, speech recognition and machine translation are now being successfully applied to problems in regulatory genomics. However, deep learning architectures used thus far in genomics are often directly ported from computer vision and natural language processing applications with few, if any, domain-specific modifications. In double-stranded DNA, the same pattern may appear identically on one strand and its reverse complement due to complementary base pairing. Here, we show that conventional deep learning models that do not explicitly model this property can produce substantially different predictions on forward and reverse-complement versions of the same DNA sequence. We present four new convolutional neural network layers that leverage the reverse-complement property of genomic DNA sequence by sharing parameters between forward and reverse-complement representations in the model. These layers guarantee that forward and reverse-complement sequences produce identical predictions within numerical precision. Using experiments on simulated and *in vivo* transcription factor binding data, we show that our proposed architectures lead to improved performance, faster learning and cleaner internal representations compared to conventional architectures trained on the same data.

**Availability:** Our implementation is available at https://github.com/kundajelab/keras/tree/keras_1

## 1 Introduction

Deep learning models such convolutional neural networks have been recently used to decipher predictive regulatory patterns encoded in genomic DNA sequence [1] [5] [8]. Transcription factor (TF) complexes bind combinations of sequence motifs in non-coding regulatory elements to regulate gene expression. While sequence affinity models of individual transcription factors are reasonably well characterized, the complex combinatorial sequence grammars that specify *in vivo* transcription factor binding and define cell type-specific regulatory elements remain largely uncharacterized. Deep learning models are particularly enticing for this problem because they are capable of inducing hierarchical, predictive patterns of increasing complexity from raw input DNA sequences without relying on explicit featurization (such as featurization into *k*-mers). Deep convolutional neural networks (CNNs) consist of multiple, hierarchical layers of elemental pattern matching units known as convolutional filters that can learn distributed representations of predictive sequence motifs and grammars from labeled sets of raw DNA sequences (e.g. sequences bound or unbound by a TF based on an *in vitro* or *in vivo* TF binding assay). Each convolutional filter encodes a predictive sequence pattern, akin to position weight matrices (PWMs), in the form of a 2D matrix of weight parameters with 4 rows representing the 4 nucleotides and *l* columns representing consecutive positions of the pattern. An input DNA sequence is scanned (convolved) against each convolutional filter providing a sequence of positional pattern matching scores. Convolutional filters in subsequent layers operate collectively on the positional match scores from all filters from the previous layer, thereby encoding higher-order patterns.

However, popular CNN architectures derived from other application domains such as computer vision do not take advantage of properties of data modalities specific to the genomics. In double-stranded DNA, the same pattern may appear identically on one strand and its reverse (in opposite orientation) due to complementary base pairing. i.e. a sequence pattern (e.g. ACG) observed on one strand is equivalent to its reverse-complement (RC) pattern (e.g CGT) on the same strand. In traditional CNN architectures, a pattern and its reverse complement would be learned as separate convolutional filters or artifactual palindromic filters, thereby resulting in inefficient use of weight parameters, reduced support for each pattern and its RC and more unstable models. Without explicit weight constraints, these filters may contribute very differently to the final prediction of the entire network resulting in wildly different predictions for forward and RCs of the same sequence. This disparate treatment represents a biological inconsistency of current deep learning models for genomics.

We develop reverse-complement (RC) convolutional neural networks to enable biologically valid learning in which forward and reverse-complement sequences are treated identically [**Fig. 1**]. First, we implement RC convolutional filters that share weights between forward and RC patterns. We then developed RC batch normalization, RC weighted sum layers and RC dense layers to preserve the reverse complement weight-sharing through all layers of the network leading to the final predictions (See **Section 3.1**). We demonstrate that using a RC architecture can lead to higher predictive performance of models trained on simulated and *in vivo* TF binding datasets to discriminate between bound and unbound sequences. RC architectures also learn cleaner internal representations. These improvements can be particularly useful with limited training data involving TFs with a few hundred bound sequences. Most importantly, the networks have a biologically consistent interpretation for forward and RC sequences.

**Fig. 1.**
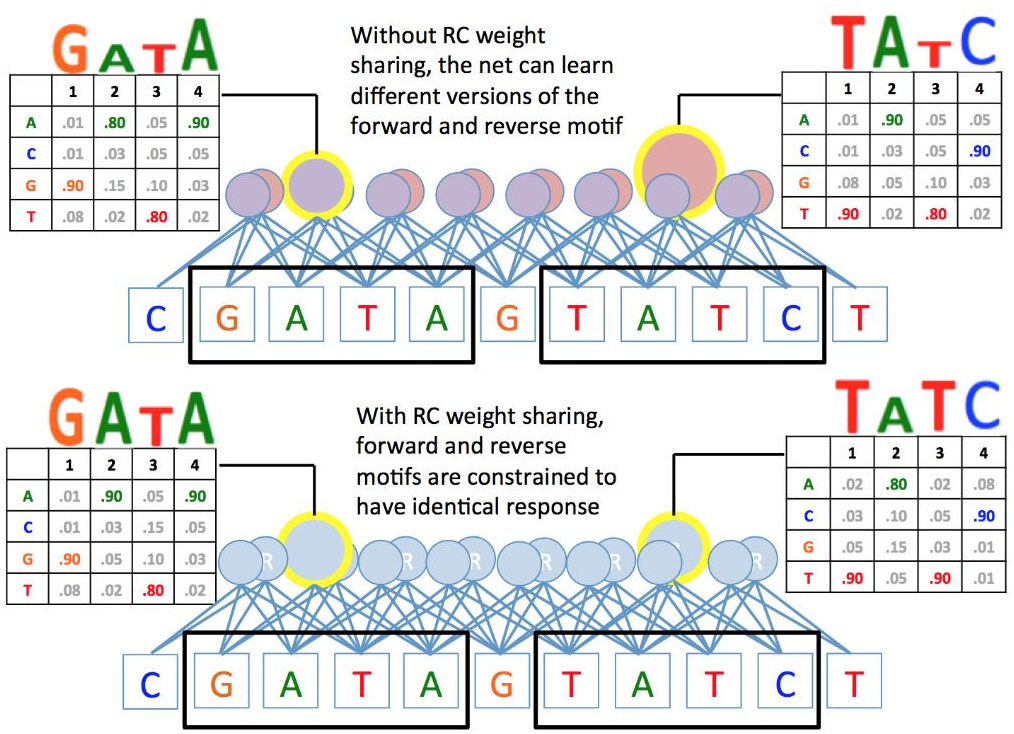
Illustration of the core idea behind reverse-complement weight sharing. In traditional architectures, the network learns completely separate representations for forward and reverse-complement versions of patterns. By using reverse-complement weight sharing, we can constrain these representations to be consistent with each other. As a result, the model gives identical responses when it sees forward and reverse-complement versions of a pattern.

## 2 Results

### 2.1 Embedded motif simulations

First, we compare performance of traditional and RC architectures on simulated TF binding prediction tasks. We sample motif instances from the canonical PWMs of 3 TFs (GATA1, ELF1 and RXRA) and randomly embed the sampled motif instances into 10K random sequences of length 200 bp for each TF (30K sequences in total) such that motif instances of a TF may appear 1-3 times in each sequence (See **Section 3.2**). We shuffle 20% of the labels to increase the difficulty of learning. We set up three separate binary classification tasks to discriminate sequences containing motifs of each TF against sequences containing motifs of the other two TFs. We train multi-task convolutional neural networks to jointly learn all three binary prediction tasks.

We compare three architectures: a RC model with 20 filters in the convolutional layer and traditional models with 20 or 40 filters in the convolutional layer (See **Section 3.4**). Note that while the traditional model with 20 filters has roughly the same number of parameters as the RC model with 20 filters (See **Section 3.4.1**), the traditional model with 40 filters consumes roughly the same amount of memory (RAM) as the RC model with 20 filters. To demonstrate robust improvements, we re-train the models with 10 different random seeds and compare the validation set auROCs and auPRCs (averaged across the three tasks).

#### 2.1.1 Reverse-complement weight sharing improves validation set performance

The RC architecture consistently outperforms both traditional architectures across all 10 random seeds [**Fig. 2, Fig. 3**]. We note that the RC architecture with 20 filters per layer achieves superior validation set performance compared to a traditional architecture with 20 filters per layer despite having similar training set performance. By contrast, the traditional model with 40 filters per layer gives no benefit to validation set performance despite achieving higher training set performance. Thus we note both an improvement in validation performance as well as a reduction in overfitting by using a RC architecture.

**Fig. 2.**
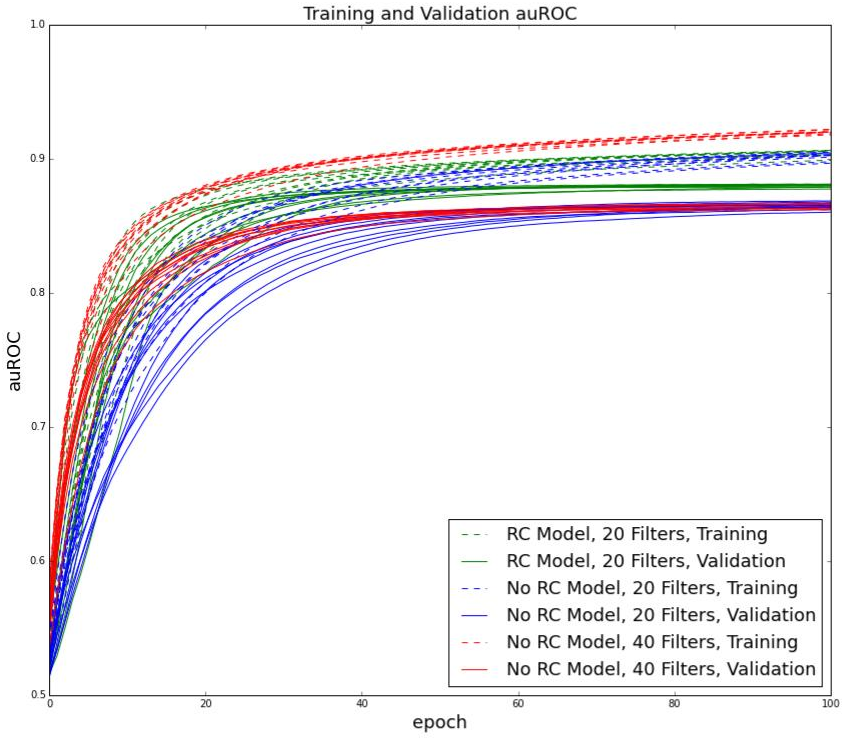
Architecture with RC weight sharing improves validation set performancewhile maintaining training set performance. We plot the validation set (solid line) and training set (dotted line) auROCs for the first 100 epochs across all 10 uniquely seeded models for each type of architecture. Note that the traditional model with twice the number of filters per layer shows higher training set performance with no corresponding improvement in validation set performance.

**Fig. 3.**
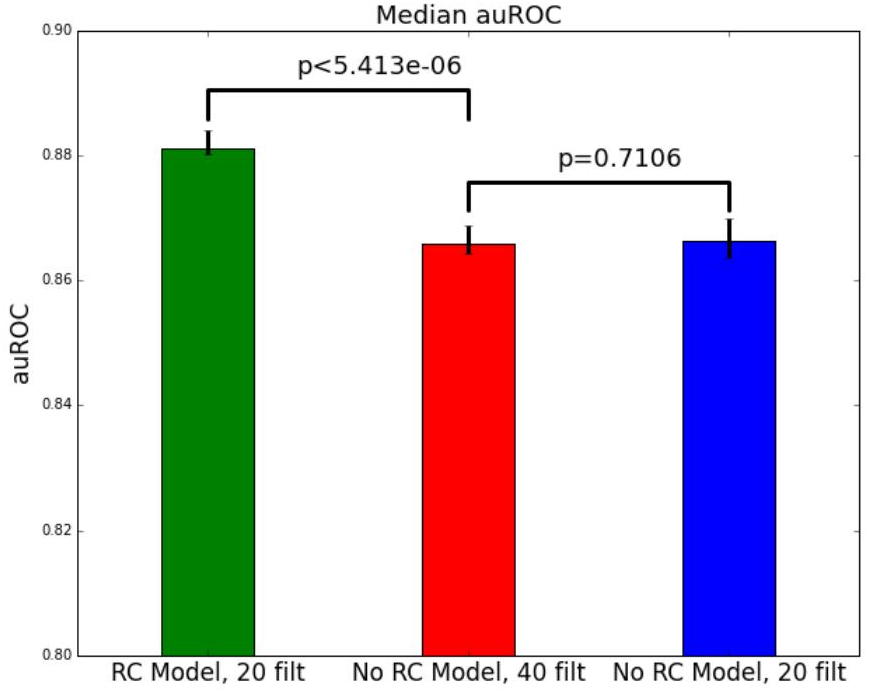
The RC architecture improves performance over traditional architectures on simulated data. Error bars show min and max performance over 10 random seeds. The RC architecture is significantly better (p<5.413e-06) than both traditional architectures. The traditional architecture with 40 filters does not show significantly different validation set performance compared to the traditional architecture with 20 filters (p=0.7106). P-values were computed using a one-sided exact Wilcoxon test (see Section 3.8).

#### 2.1.2 Reverse-complement weight sharing enables more informative updates in stochastic gradient descent

We note that the learning curves in [**Fig. 2**] for the RC architecture with 20 filters per layer consistently lie above the learning curves for the traditional architecture with 20 filters per layer, confirming our intuition that adding RC weight sharing enables the model to make more informative updates per batch. Note however that this does not necessarily translate into a reduction in wall-clock time, as the RC architecture has a larger number of output channels (and therefore performs more computations) than the equivalent traditional architecture with the same number of filters. We also note that the learning curves for the traditional architecture with 40 filters per layer are initially higher, implying that adding more parameters to the model may enable it to learn more quickly during initial training epochs even if it does not benefit final performance. See the Discussion for more details.

### 2.2 Transcription Factor binding data

Next, we evaluated relative performance of RC and traditional architectures on *in vivo* TF binding prediction tasks using chromatin immunoprecipitation sequencing (ChIP-seq) datasets for CTCF, SPI1 and MAX in the GM12878 cell-line. We set up binary classification tasks where the positive set consisted of sequences underlying high-confidence ChIP-seq peaks for a given TF, and the negative set consisted of chromatin accessible DNase peaks in GM12878 that were not bound by the TF. All models had 3 convolutional layers and a logistic output neuron except for the models used in [**Fig. 5**], which had 1 convolutional layer.

#### 2.2.1 Reverse-complement weight sharing guarantees consistency between forward and reverse-complement versions of a sequence

Ideally, forward and RC versions of a sequence should produce identical predictions from a model in the absence of any strand-specific properties of the training data. Unfortunately, traditional deep learning models do not enforce this constraint and can produce wildly different predictions on forward and RC sequences [**Fig. 4a**]. While augmenting the training set with the RCs of all the original sequences can alleviate the problem [**Fig. 4b**], it far from eradicates it. In contrast, the architectures with RC weight sharing produce identical predictions on forward and RC sequences by design [**Fig. 4c**].

**Fig. 4.**
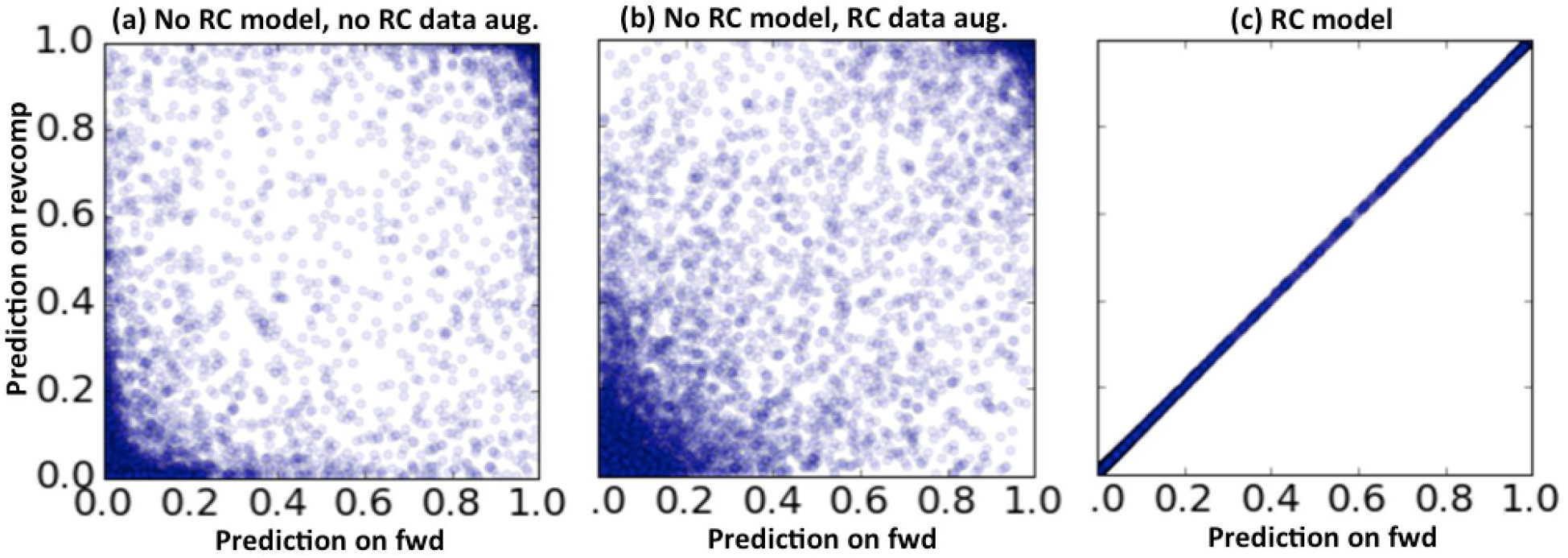
RC weight sharing guarantees identical predictions on forward and RC versions of a sequence within numerical precision. Models were trained on chromosome 1 ofGm12878 CTCF ChIP-seq data (see Sections 3.3 and 3.4.2). Scatter plots show consistency of predictions on forward and RC inputs for 10,000 sequences from the testing set. X-axis is model output on the original sequence and y-axis is model output on the RC. **(a)** Model with 16 filters per layer trained without RC weight sharing and no data augmentation **(b)** Model with 16 filters per layer trained without RC weight sharing but with the training set augmented with RCs **(c)** Model with 16 filters per layer trained with RC weight sharing.

#### 2.2.2 Reverse-complement weight sharing leads to cleaner internal representations

To understand how RC weight sharing influenced the internal representations learned by the models, we trained 1-layer models with and without RC weight sharing on chr21 of the Gm12878 CTCF ChIP-seq data. We used 8 convolutional filters for the model with RC weight sharing and 16 filters for the traditional model. We found that the model with RC weight sharing was able to learn a filter that strongly resembles the canonical CTCF motif, while the model lacking weight sharing showed a tendency to collapse the forward and RC versions of the CTCF motif into “palindromic” representations [**Fig. 5**]. We hypothesize that this is because the palindromic representation is a local minimum in the optimization; without RC weight sharing, a single filter cannot simultaneously represent high-fidelity versions of the forward and RC motifs, and thus gradient descent might cause filters to gravitate towards “hybrid” representations that do a modest job of representing both at once. We confirmed that the trends illustrated in **Fig. 5** were representative by repeating the filter visualization on models trained with multiple different random seeds. The poor representations learned by the model lacking RC weight sharing were not caused by having twice the number of filters per layer; traditional models that used only 8 convolutional filters (rather than 16) tended to learned a single palindromic filter, with the remaining filters contributing little to none to the prediction. See **Sections 3.3 3.4.2 and 3.5** for details.

**Fig. 5.**
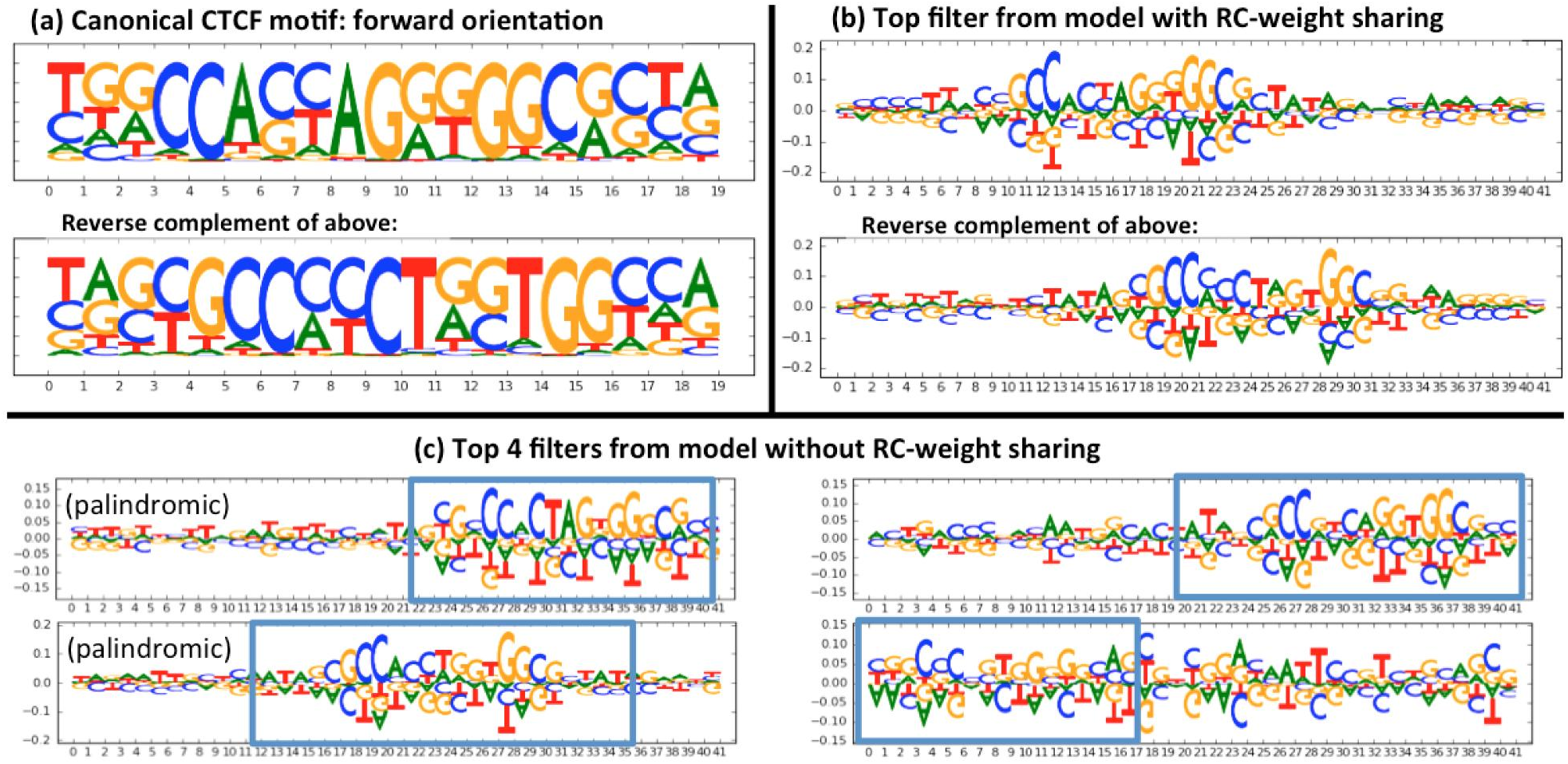
RC weight sharing leads to cleaner internal representations. Weights were visualized for simple 1-layer models trained on chromosome 21 of the Gm12878 CTCF datawith and without RC weight sharing. Details of model architecture and training are described in Sections 3.4.2. **(a)** Canonical “CTCF_known1” motif from ENCODE [6] and reverse complement. **(b)** Top filter (out of 8) from the model trained with RC weight sharing. **(c)** Top 4 filters (out of 16) from the model trained without RC weight sharing - note that the top 2 filters learned “palindromic” patterns that are RCs of themselves. Also note that the true CTCF motif is not palindromic. Any filters not depicted were not found to contribute substantially in their respective models.

#### 2.2.3 Reverse-complement weight sharing leads to slower degradation in performance with decrease in training set size

To test the impact of training set size, we trained models on 5%, 10%, 20%, 40%, 60% and 80% of the regions on chr1 of the Gm12878 CTCF ChIP-seq data set and observed the degradation in validation set performance with training set size [**Fig. 6**]. We compared a model with RC weight sharing that had 16 convolutional filters per layer to models without RC weight sharing that had 16 or 32 convolutional filters per layer. See **Sections 3.3 and 3.4.2** for details. We find that the model with RC weight sharing displays consistently higher auROC and auPRC and performance appears to drops off more slowly with decreasing training set size compared to the other two models. As a reminder, a model with 16 filters per layer and RC weight sharing has fewer parameters but consumes roughly the same amount of RAM compared to a model with 32 filters per layer and no RC weight sharing.

**Fig. 6.**
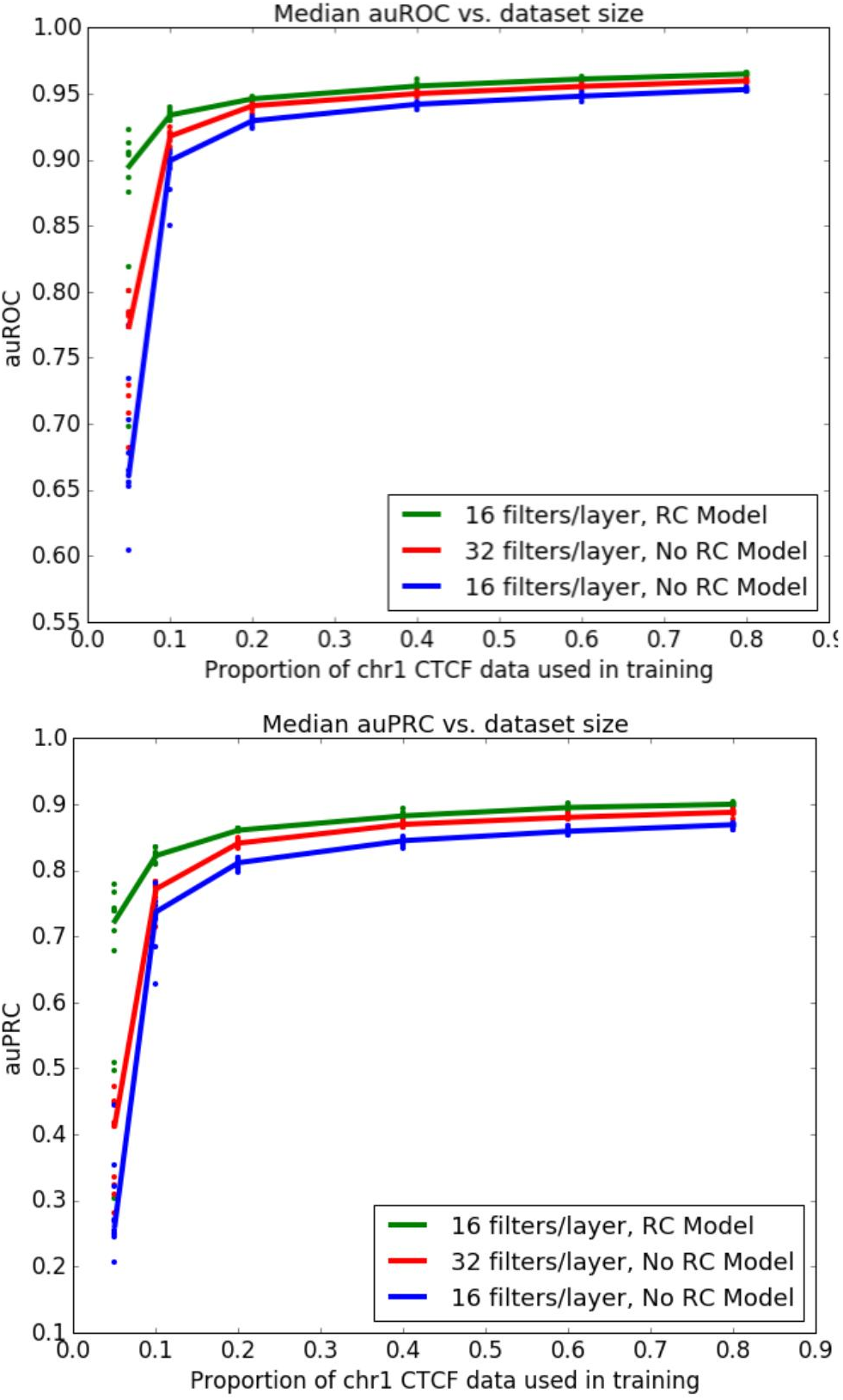
RC weight sharing leads to slower degradation in performance with training set size. Shown are the median auROC and auPRC on the validation set across 10 random seeds at different training set sizes (each dot represents a different random seed). Green line represents a model with RC weight sharing. Blue line represents a model without RC weight sharing that has the same number of filters per layer as the green line. Red line represents a model without RC weight sharing that has twice the number of filters per layer as the green line and therefore consumes roughly the same amount of RAM.

#### 2.2.4 Reverse-complement weight sharing leads to superior performance

Consistent with the results on simulated data, we found that models using RC weight sharing gave superior performance on the validation set compared to traditional models with the same or double the number of filters per layer [**Fig. 7**]. As a reminder, traditional models with twice the number of filters per layer consume roughly the same amount of RAM as models using RC weight sharing. Our results were robust to changes in the number of filters [**Fig. 7a**] as well as across the different TFs with diverse binding landscapes and sequence affinities. [**Fig. 7b**].

**Fig. 7.**
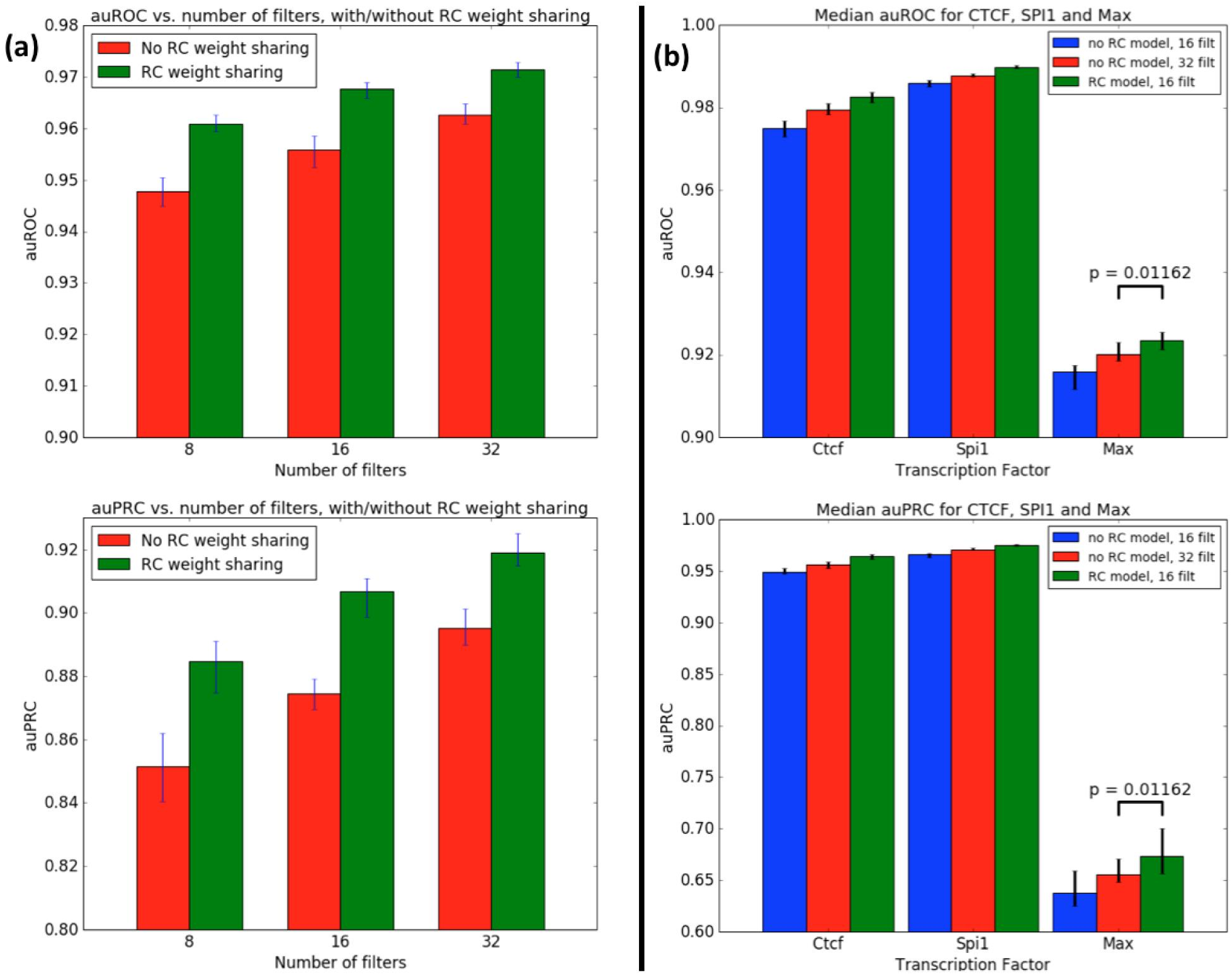
RC weight sharing improves performance over traditional architectures on ChIP-seq data. We trained traditional models and RC models with different numbers of filters per layer and on different TF binding data sets. a Validation set auROC and auPRC of RC models and traditional models with 8, 16 and 32 filters per layer trained on chr1 of the CTCF data set. We subset the training set to chr1 because CTCF has a very large number of peaks and we wanted to show behavior on a training set of a size that was more typical of most TFs. RC models show a consistent improvement over traditional models with the same or double the number of filters per layer b Validation set auROC and auPRC of RC models and traditional models with 16 filters per layer trained on the Gm12878 TF binding data for CTCF, SPI1 and MAX. Here, we used the full training set for all TFs. The performance of a traditional model with 32 filters per layer is also shown for comparison. Error bars show min and max performance over 10 different random seeds. p-value was computed using a 1-sided exact Wilcoxon test. See Section 3.8 for details.

## 3 Methods

### 3.1 Layers adapted for reverse-complement weight sharing

We describe the four novel layers that enable complete weight sharing between forward and RC representations of the input.

#### 3.1.1 Reverse-Complement Convolution

Consider the simple case of a filter in the first convolutional layer operating on one-hot encoded ACGT sequence input. We can compute the reverse complement of this filter by first reversing the weights in the length dimension and then exchanging the weights of A with T and C with G and vice versa. Note that if the input channels were ordered as ACGT, this amounts to simultaneously reversing the weights in both the length and channel dimensions.

Now consider the case of a filter operating in a later convolutional layer which takes as its input the output of some previous convolutional layer. How can we find the RC of this filter? If we do not have any information about the input channels, computing the RC is difficult to do. However, if we are told that the input channels *i* and *n* – 1 – *i* are RCs of each other (that is, the input channel at index *i* detects the reverse complement of the pattern detected by the input channel at index *n* – 1 – *i*, where n is the number of input channels), we can once again compute the RC by inverting the weights in both the length and channel dimensions. For this reason, when implementing our RC layers, we always maintain the property that the output channel at index *i* represents the RC of the output channel at index *n* – 1 – *i* (where n is the number of output channels); it enables us to compute the RC in downstream layers by doing simple inversions of the weight matrices. This is illustrated in [Fig. 8]. For a given number of input channels, filter length, and number of filters, the RC convolutional layer has the same number of parameters as a standard convolutional layer. However, it has twice the number of output channels and therefore uses twice the RAM. Our current implementation is for a 1D convolution, but is easily extendable to implementations using additional dimensions.

**Fig. 8.**
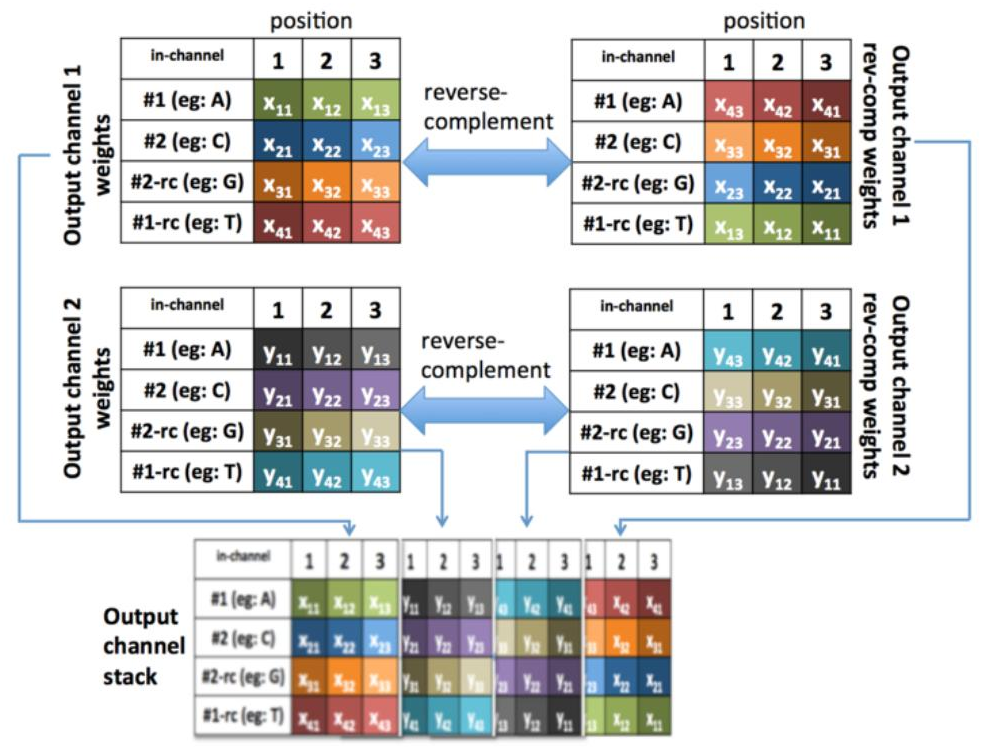
Weight sharing in a RC convolutional layer. We depict two filters of size [4, 3] thatwill scan an input with 4 channels. The input channels may be one-hot encoded ACGT (as is the case for the first convolutional layer) or may be the output of a previous RC convolutional layer. The only constraint on the input is that the RC of the channel at index *i* is present at index *n* – 1 – *i* (where *n* is the number of input channels; note that ACGT satisfies this constraint). The two output filters are individually reverse complemented (length and channel axes are inverted) and then appended as additional channels in the opposite order so as to maintain the property that the reverse complement of the channel at index *i* is present at *n* – 1 – *i* (where *n* is the number of output channels). This structure allows us to compute reverse complements in downstream layers by simply inverting the length and channel dimensions.

#### 3.1.2 Batch Normalization for Reverse-Complement Convolutions

Batch normalization for standard convolutional layers computes normalization parameters on a per-channel basis; that is, the activations for all neurons in a given channel are used to compute the batch-statistics for the channel, regardless of the position in the length dimension. We can leverage this implementation detail to compute batch statistics for all neurons in both the forward and RC of a channel as follows: we first rearrange the incoming tensor so that the activations of the RC channels are concatenated along the length dimension to their forward counterparts (as a result, we halve the size of the tensor along the channel dimension and double it along the length dimension). We then feed this rearranged tensor to the batch-normalization calculation for standard convolutional layers, in the process updating α and β for the halved set of input channels. Then, we take the normalized tensor and rearrange it so that the RC activations are returned to their original positions in the incoming tensor.

#### 3.1.3 Reverse-Complement Weighted Sum

In the standard convolutional architecture, a pooling layer feeds into a fully connected layer. This fully connected layer treats every neuron in the pooling layer completely independently, without any regard to its position in the channel or length dimensions. Consider the case where each channel of the pooling layer has a positional pattern - for instance, if a channel represents the presence of a motif, that motif may preferentially occur towards the center of the region rather than the flanks; in this situation, the net should learn to upweight motif instances that occur in the center. One undesirable aspect of going from a pooling layer straight to a fully-connected layer is that every neuron in the fully-connected layer must learn such positional patterns completely independently. A strategy for alleviating this is related to the idea of a separable convolution [10]; we can learn a positional weighting for each channel separately and then combine the positionally-weighted channel outputs. To this end, we developed the Weighted Sum layer, which was implemented as follows:

Assume the input is a two-dimensional matrix of size (input_length × num_channels), where *x_ic_* represents the input at position *i* and channel *c*. The Weighted Sum layer has a weight matrix also of size (input_length × num_channels), where *w_ic_* is the weight for channel *c* at position *i*. The output of the weighted sum layer is a vector of length (num_channels), where *y_c_* (the output for channel *c*) is:

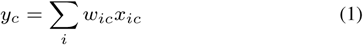

How can we now incorporate RC information? If we make the assumption that the input channel at index *c* is the RC of the input channel at index (*n* – 1 – *c*) (where *n* is the number of input channels), we can expect that the RC channel shows the reverse positional properties of the forward channel (and if the positional properties are symmetric along the length axis, we can expect the RC channel to show identical positional properties as the forward channel). Thus, we have the constraint that *w_ic_ = w_(l–i)(n–1–c)_*, where *l* is the input length and *n* is the number of input channels. We can achieve this constraint by reversing the positional weights for the forward channels and concatenating them in the opposite order to get the weights for our RC channels. Note that in our experiments, the Weighted Sum layer did not use bias terms and was not followed by a nonlinearity.

When developing the Weighted Sum layer, we found that the default Keras initialization strategy of glorot_uniform did not work well. We ultimately decided to initialize the layer to match the corresponding weight distribution in the architecture lacking the Weighted Sum layer. This initialization is implemented in our code as “fanintimesfanouttimestwo” (see **Section 3.7**); we find that it works well but can likely be further improved (see Discussion).

#### 3.1.4 Dense Layer after Reverse-Complement Weighted Sum

How do we design a Dense layer after a Weighted Sum layer to leverage RC information? We note that if the Weighted Sum layer has RC weight sharing enabled, then the channel at index *i* is the reverse complement of the input at index *n* – 1 – *i*, where n is the total number of channels. If we assume that forward and RC signals are functionally equivalent at this layer, it stands to reason that if the output neuron at index *j* puts a weight of *w_ij_* on the input channel at index *i*, it should put an identical weight on the input channel at index (*n* – 1 – *i*). Thus, for every neuron in the Dense layer, we can obtain the weights on the RC channels by using the weights on the corresponding forward channels. When implemented, this amounts to reversing the top half of the weight-matrix along the input dimension and concatenating it to form the bottom half.

Note that forward and RC representations converge at this layer - that is, assuming the other RC weight sharing layers were used where appropriate earlier in the network, forward and RC sequences are guaranteed to produce identical activations at this layer within numerical precision. Even if the network has more downstream layers that follow this layer, no additional RC weight sharing is necessary to maintain this guarantee. An illustration of a network with complete RC weight sharing is provided in [**Fig. 9**].

**Fig. 9.**
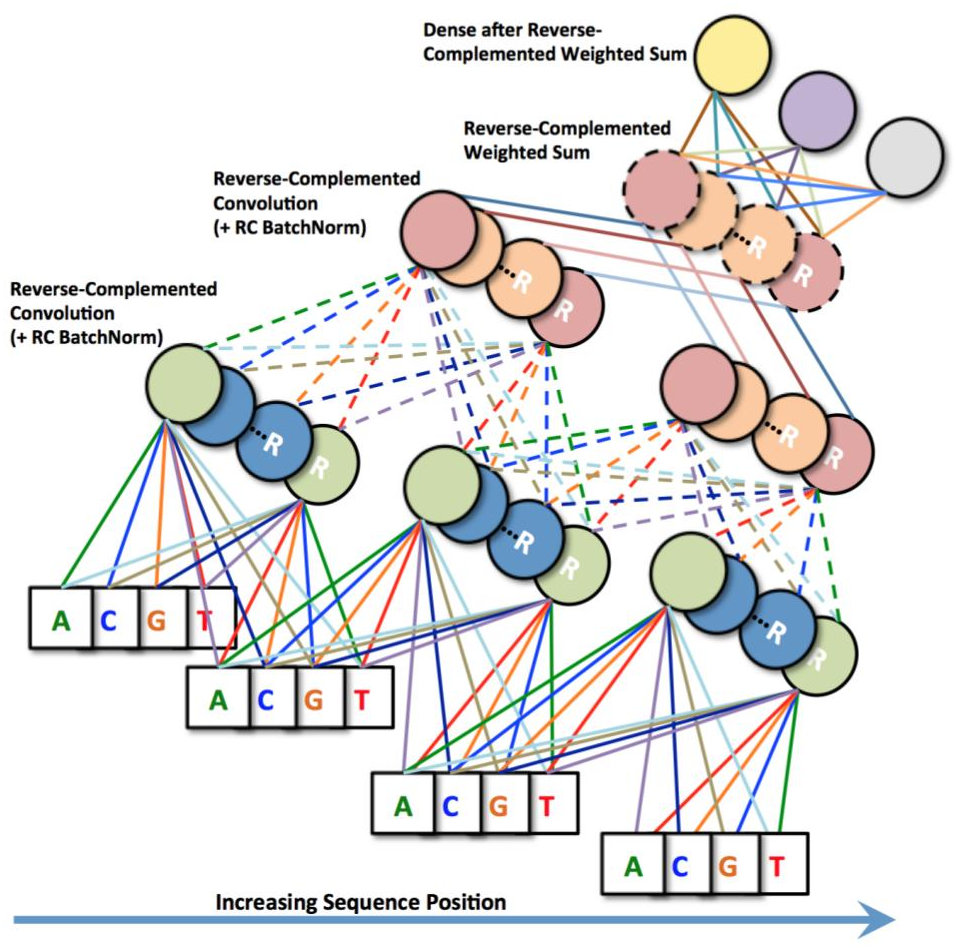
Complete RC architecture. We illustrate two RC convolutional layers (with optional batch normalization) scanning an input sequence with filter width 2. These layers are followed by a Weighted Sum layer and a final Dense layer, both of which have RC weight sharing. Edges that have the same color and style represent identical weights. Neurons with the same interior color and border have shared weights, with the R’s denoting RC versions as explained in Fig. 8.

### 3.2 Simulated datasets

We used the GATA_known6, ELF1_known2, and RXRA_known1 Position Weight Matrices (PWMs) from the ENCODE consortium [6] and simulated a total of 30,000 sequences (10,000 per TF). Sequences were simulated by (1) generating a random background sequence of length 200bp with 40% GC content, (2) determining the number of instances of the motif to insert by sampling from a Poisson distribution, (3) sampling the motif instances determined in (2) from the PWM of the motif, (4) reverse complementing each sampled instance with probability 0.5, and (5) inserting each sampled instance at a random position within the sequence with the constraint that it does not overlap a previously inserted instance. In step (2), the Poisson distribution had a mean of 2 but was truncated to have a max of 3 and a min of 1 (if a number was sampled that was greater than 3 or smaller than 1, it was resampled from the Poisson until it was in the appropriate range). We randomly allocated 30% of the data to validation, 30% to testing, and 40% to training. We also shuffled 20% of the labels to simulate mislabeling thereby increase the difficulty of the learning task.

### 3.3 ChIP-seq datasets

TF ChIP-seq datasets for Gm12878 were obtained from the ENCODE consortium [3]. For CTCF, we used the specific file “wgEncodeBroadHistoneGm12878CtcfStdAlnRep0”. For SPI1, we used “wgEncodeHaibTfbsGm12878Pu1Pcr1xAlnRep0” and for MAX we used “wgEncodeSydhTfbsGm12878MaxIggmusAlnRep0”. Preparation of the positive and negative sets for a give TF was as follows: for the positive set, we used 1000bp windows centered around the summits of rank-reproducible peaks [9]. For the negative set, we identified those DNase peaks in Gm12878 that did not overlap the top 150K relaxed ChIP-seq peaks of the TF (where the “relaxed” peaks were called by SPP at a 90% FDR) and used 1000bp centered around their summits. The resulting sizes of the positive and negative sets were as follows:

For experiments involving chr1 for CTCF and any subsets, the training set contained regions on chr1 (or some subset), the validation set contained regions on chr17 and the test set contained regions on all other chromosomes. For experiments involving chr21 for CTCF, the training set consisted of regions on chr21, the validation set consisted of regions on chr22, and the test set contained regions from all other chromosomes. For [**Fig. 7b**], the training set consisted of all regions except regions on chr1 and chr2, the validation set consisted of regions on chr1, and the testing set consisted of regions on chr2.

### 3.4 Model Architectures

We trained models using Keras [2] with the Theano backend [12]. All references to convolutional layers are for 1D convolutions. The input sequences had ACGT one-hot encoded along the channel dimension.

#### 3.4.1 Models trained on simulated data

All models trained on simulated data had one convolutional layer with filter width 21 and stride 1, followed by batch normalization [4] and a ReLU nonlinearity. The number of filters in the convolutional layer was varied according to the experiment; the architecture with RC weight sharing used 20 filters, and the traditional architecture used 20 or 40 filters. The ReLU nonlinearity was followed by a max-pooling layer of width and stride 20. In the case of RC weight sharing, the max-pooling layer was followed by a Weighted Sum layer which then fed into a RC dense layer with three sigmoid output neurons (one for each of the three motif tasks). In the case of no RC weight sharing, the max-pooling layer fed directly into the dense layer. The models using RC weight sharing applied this weight sharing at all appropriate layers (convolutional, batch normalization, weighted sum and dense layers). Models were trained using the Adam optimizer [7] with a learning rate of 0.001 and a binary cross-entropy loss. Training was terminated when the mean validation set auROC across all 3 tasks showed no improvement after 10 epochs, where each epoch consisted of 5,000 samples in batches of 500. Training and validation set auROC and auPRC (averaged across all 3 tasks) were recorded at the end of each epoch. The order of the training examples was randomly shuffled before training commenced. All experiments were repeated across 10 random seeds to assess the robustness of the results.

Note that the traditional model with 20 filters has roughly the same number of parameters as the RC model with 20 filters: the traditional model has 20 *(4 * 21 + 1) parameters in the convolutional layer, 20 * 2 parameters in batch normalization layer and ((180/20) * 20 + 1)* 3) parameters in the fully-connected layer for a total of 2; 283 parameters, while the RC model has the same number of parameters in the convolutional layer, 40 2=2 parameters in the batch normalization layer, (180=20) 40=2 parameters in the weighted sum layer and ((40/2) + 1)* 3 parameters in the dense layer for a total of 1; 983 parameters. Intuitively, this is because although the output size of the convolution is doubled, the number of parameters in subsequent layers is halved, leading them to approximately cancel out except for a small difference introduced by the Weighted Sum layer. Refer to **Section 3.6** on “Calculation of the number of parameters” for a breakdown of how to do the calculation for each layer.

#### 3.4.2 Models trained on ChIP-seq data

Except for the models trained to visualize the learned filters (described in the next section), all models trained on ChIP-seq data used a network with 3 convolutional layers of stride 1 and widths 15, 14 and 14 respectively (for sequences of length 1000, this means that the output size of the third convolution is of length 960). The number of filters per convolutional layer was varied according to the experiment. Every convolutional layerused batch normalization [4] followed by a ReLU nonlinearity. The last convolutional layer fed into a max-pooling layer that had pooling width 40 and stride 20 (there was no max-pooling between convolutional layers). In the case of no reverse-complement weight sharing, the max-pooling layer fed into a single sigmoid output neuron. In the case of RC weight sharing, the max-pooling layer was followed by a Weighted Sum layer, which then fed into a single sigmoid output neuron. The models using RC weight sharing applied this weight sharing at all appropriate layers (convolutional, batch normalization, weighted sum and dense layers). For the CTCF, SPI1 and MAX models, class weights of 5:1, 5:1 and 16:5:1 were used to upweight the positive examples, which roughly matched the class imbalance in our dataset (see Table 1). Models were trained with the Adam optimizer [7] with a learning rate of 0.001 and binary cross-entropy loss. Models were trained for 80 epochs, where each epoch consisted of showing the model 5,000 samples in batches of 100. Validation set performance was evaluated at the end of every epoch, and the best validation set performance over the 80 epochs as well as the corresponding model were recorded. The order of the training examples was randomly shuffled before training commenced. All experiments were repeated across 10 random seeds.

**Table 1.** Number of positive and negative examples after data processing

| TF | Positives | Negatives | Negatives:Positives |
| --- | --- | --- | --- |
| CTFC | 44,982 | 225,533 | 5.01 : 1 |
| SPI1 | 42,938 | 203,960 | 4.75 : 1 |
| MAX | 12,542 | 206,628 | 16.47 : 1 |

When there is a single output neuron, the weights connecting the Weighted Sum layer to the Dense layer are redundant, as they effectively scale the weights of the preceding Weighted Sum layer. For this reason, we froze the weights of the sigmoid output neuron to be a vector of all 1’s. This was to ensure that any differences we observed were due to RC weight sharing and not due to differences in the behavior of the optimizer caused by introducing redundant parameters. We discuss the initialization of the Weighted Sum layer in **Sections 3.7**.

### 3.5 Visualizing the learned representations

For the models depicted in [**Fig. 5**], we used a similar setup to what was used in the other ChIP-seq experiments (described above), but included 1 convolutional layer (with batch normalization) instead of 3. The RC model depicted had 8 filters in the convolutional layer and the traditional model had 16 filters. The training data for the traditional model was augmented with reverse-complementation.

Contribution scores in the convolutional layer were scored by computing the gradient of the logit of the sigmoid w.r.t. a neuron in the convolutional layer and multiplying the gradient by the activation of the neuron. This can be interpreted as a Taylor approximation of the change in the logit if the filter activation is set to 0 [11]. The score of a particular filter for a given sequence was found by summing the contribution scores across all positions in the sequence. A final score for each filter was then found by subtracting the average filter score across negative examples from the average filter score across positive examples. Most filters had a score of around zero, but a handful of filters had notable positive scores; these filters are depicted in [**Fig. 5b**] and [**Fig. 5c**]. Filter weights were mean-centered prior to visualization using the output-preserving transformation described in **Section 2.6** of Shrikumar et al. [11].

### 3.6 Calculation of the number of parameters

We explain here how to calculate the number of parameters in our layers.

#### 3.6.1 Convolutional layers

For both standard convolutions and convolutions with RC weight sharing, the number of weight parameters is (number_of_input_channels × filter_length × number_of_output_filters), and the number of bias parameters is (number_of_output_filters). However, note that for a standard convolutional layer the number of output channels is the same as the number of output filters, whereas for convolutional layers with RC weight sharing the number of output channels is twice the number of output filters (because the output channels include the results of applying both the forward and RC versions of a filter).

#### 3.6.2 Batch Normalization layers

For a standard batch normalization layer following a convolutional layer, the number of parameters is (2 × number_of_input_channels), with one set of *α* and *β* one set of parameters per channel. For a batch normalization layer designed to follow a convolutional layer with RC weight sharing, the number of parameters is cut in half because the *γ* and *β* parameters are shared between the forward and RC channels.

#### 3.6.3 Length of output of max-pooling layer

To calculate the number of parameters going into a Dense layer or Weighted Sum layer, it is necessary to calculate the length of the output of the max-pooling layer. Every convolutional layer of stride 1 reduces the length of the output by (filter_width – 1). The output length of the max-pooling layer is then found as 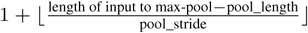

#### 3.6.4 Weighted Sum layers

Without RC weight sharing, a weighted sum layer would have a number of parameters equal to (number_of_input_channels × length_of_previous_layer), where the previous layer in our case is a max-pooling layer. With RC weight sharing, the number of parameters is cut in half as parameters are shared between the forward and RC channels.

#### 3.6.5 Dense layers

Without RC weight sharing, a dense layer has (size_of_input × size_of_output) weight parameters and (size_of_output) bias parameters. Note that when a dense layer follows a max-pooling layer, the size of the input is equal to (length_of_pooling_layer × number_of_channels). For dense layers designed to follow a Weighted Sum layer with RC weight sharing, the number of weight parameters is cut in half as the weight on a given channel is tied to the weight on its RC counterpart.

### 3.7 Initialization of the Weighted Sum layer

The default initialization mode implemented in Keras is “glorot_uniform”. Briefly, glorot_uniform initializes the weights to be drawn from a uniform distribution with a min of *–s* and max of *s*, where *s* is computed as:

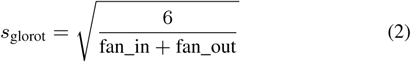

In Keras, fan_in and fan_out are computed according to the shape of the weight matrix. When the weight matrix is two-dimensional (as is the case for dense layers), fan_in is the length of the first dimension of the matrix (which corresponds to the number of input neurons per output neuron) and fan_out is the length of the second dimension (which corresponds to the number of output neurons).

Consider a max-pooling layer that has *c* channels and length *l* (here, *c* includes any RC channels). When this max-pooling layer is followed by a dense layer, the input dimension has size *c* × *l*. Thus, fan_in = *c* × *l*. Assuming that the number of output neurons (fan_out) is small relative to *c* × *l*, we find that *s*_glorot_ is approximately 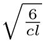

How can we obtain a comparable distribution of weights in the Weighted Sum layer that uses RC weight sharing? The weight matrix of the Weighted Sum layer has dimensions *l* × 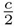, because it learns the weights at each position for each channel, and only learn the weights for the forward channels (the weights for the reverse channels are found by reverse-complementation; hence the 2 in the denominator). Keras would thus compute fan_in for this weight matrix to be *l* and fan_out to be 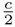. If we use an initializer that computes s according to the following formula:

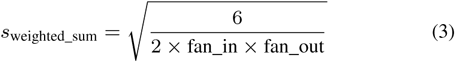

Then when we substitute the values for fan_in and fan_out computed from the weighted sum layer, we get 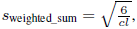 as desired.

### 3.8 Statistical tests

All *p*-values were calculated using the one-sided Wilcoxon test implemented in R (exact calculation, i.e. no normal approximation). Values came from running each experiment with 10 different random seeds.

### 3.9 Implementations

A gist illustrating model setup is at: https://gist.github.com/AvantiShri/21ec8de292b9c0c91e4d448b5c02a118. Commit-specific permalinks to code implementations are at:

- keras.layers.convolutional. RevCompConv1D: https://github.com/kundajelab/keras/blob/2d329/keras/layers/convolutional.py#L175-L267
- keras.layers.normalization. RevCompConv1DBatchNorm: https://github.com/kundajelab/keras/blob/2d329/keras/layers/normalization.py#L161-L271
- keras.layers.convolutional. WeightedSum1D: https://github.com/kundajelab/keras/blob/2d329/keras/layers/convolutional.py#L270-L378
- keras.layers.core. DenseAfterRevcompWeightedSum: https://github.com/kundajelab/keras/blob/2d329/keras/layers/core.py#L799-L838
- keras.initializations.fanintimesfanouttimestwo: https://github.com/kundajelab/keras/blob/2d329/keras/initializations.py#L75-L78

For the most recent implementation, refer to: https://github.com/kundajelab/keras/tree/keras_1

## Discussion

### 4.1 Importance of initialization

As mentioned in **Section 3.1.3**, the default “glorot_uniform” initialization strategy did not work for the Weighted Sum layer, which inspired us to develop an alternative initialization strategy to approximately match the corresponding weight distribution in the architecture lacking the Weighted Sum layer. It is likely that a more thoughtful initialization could lead to even better performance gains. One extension would be to initialize the weights to concord with a genomic prior, such as having higher weights towards the middle of the sequence and lower weights towards the flanks.

We also note that while traditional architectures are disadvantaged in terms of having to learn forward and RC representations separately, they do allow more free parameters for the same amount of RAM, which could enable the network to find a “lucky” initialization that resembles some version of a motif more quickly. This is corroborated by the learning curves in **Fig. 2**: the red lines are ahead in the beginning even if they fall behind later on. When combined with the observation that only a small number of filters are used relative to the total number of filters available (in simple 1-layer models initialized with 8 filters, typically only 1-2 out of the 8 would contribute substantially to the prediction), this suggests that it may be fruitful to research strategies to allow a greater proportion of convolutional filters to learn.

### 4.2 Extensions of the Weighted Sum layer

In the context of genomics, a Weighted Sum layer is arguably a very good fit because it allows the net to learn the positional properties of a transcription factor independently of how the transcription factor is used for different regulatory tasks. When followed by a large fully connected layer, it can result in substantial savings in the number of parameters. In addition to our implemented RC weight sharing, another simple extension would to make weights symmetric along the length dimension. Note that our current implementation allows for symmetric weight-sharing (set symmetric=True), but we have not studied the effects of this in detail yet. Other possible extensions include adding a nonlinearity after the Weighted Sum layer, with or without a bias term.

## 5 Conclusion

We have demonstrated that traditional convolutional networks can produce drastically different predictions on forward and RC versions of the same DNA sequence. By introducing four new layers that share weights between forward and RC representations, we can enable neural networks to model the natural RC property of inputs such as DNA sequence. In experiments on simulated and *in vivo* transcription factor ChIP-Seq data, RC architectures robustly boosted performance, reduced overfitting, frequently learned faster and produced cleaner representations compared to traditional architectures. We recommend these layers for use in convolutional neural networks trained on genomic data, particularly in cases where predictive patterns may appear in small numbers. We believe this is one of the first innovations that tailors conventional neural network architectures specifically to genomics and hope that many such improvements follow.

